# SARS-CoV-2 Uses Nonstructural Protein 16 to Evade Restriction by IFIT1 and IFIT3

**DOI:** 10.1101/2022.09.26.509529

**Authors:** Craig Schindewolf, Kumari Lokugamage, Michelle N. Vu, Bryan A. Johnson, Dionna Scharton, Jessica A. Plante, Birte Kalveram, Patricia A. Crocquet-Valdes, Stephanea Sotcheff, Elizabeth Jaworski, R. Elias Alvarado, Kari Debbink, Matthew D. Daugherty, Scott C. Weaver, Andrew L. Routh, David H. Walker, Kenneth S. Plante, Vineet D. Menachery

**Affiliations:** Department of Microbiology and Immunology, University of Texas Medical Branch, Galveston, TX, USA; Institute for Human Infections and Immunity, University of Texas Medical Branch, Galveston, TX, USA; World Reference Center for Emerging Viruses and Arboviruses, University of Texas Medical Branch, Galveston, TX, USA; Department of Pathology, University of Texas Medical Branch, Galveston, TX, USA; Department of Biochemistry and Molecular Biology, University of Texas Medical Branch, Galveston, TX, USA; Institute for Translational Sciences, University of Texas Medical Branch, Galveston, TX, USA; Department of Microbiology and Immunology, Johns Hopkins University, Baltimore, MD, USA; Department of Molecular Biology, University of California, San Diego, CA, USA; Center for Biodefense and Emerging Infectious Disease, University of Texas Medical Branch, Galveston, TX, USA

## Abstract

Understanding the molecular basis of innate immune evasion by severe acute respiratory syndrome coronavirus 2 (SARS-CoV-2) is an important consideration for designing the next wave of therapeutics. Here, we investigate the role of the nonstructural protein 16 (NSP16) of SARS-CoV-2 in infection and pathogenesis. NSP16, a ribonucleoside 2’-*O* methyltransferase (MTase), catalyzes the transfer of a methyl group to mRNA as part of the capping process. Based on observations with other CoVs, we hypothesized that NSP16 2’-*O* MTase function protects SARS-CoV-2 from cap-sensing host restriction. Therefore, we engineered SARS-CoV-2 with a mutation that disrupts a conserved residue in the active site of NSP16. We subsequently show that this mutant is attenuated both *in vitro* and *in vivo*, using a hamster model of SARS-CoV-2 infection. Mechanistically, we confirm that the NSP16 mutant is more sensitive to type I interferon (IFN-I) *in vitro*. Furthermore, silencing IFIT1 or IFIT3, IFN-stimulated genes that sense a lack of 2’-*O* methylation, partially restores fitness to the NSP16 mutant. Finally, we demonstrate that sinefungin, a methyltransferase inhibitor that binds the catalytic site of NSP16, sensitizes wild-type SARS-CoV-2 to IFN-I treatment. Overall, our findings highlight the importance of SARS-CoV-2 NSP16 in evading host innate immunity and suggest a possible target for future antiviral therapies.

**Importance:** Similar to other coronaviruses, disruption of SARS-CoV-2 NSP16 function attenuates viral replication in a type I interferon-dependent manner. *In vivo*, our results show reduced disease and viral replication at late times in the hamster lung, but an earlier titer deficit for the NSP16 mutant (dNSP16) in the upper airway. In addition, our results confirm a role for IFIT1, but also demonstrate the necessity of IFIT3 in mediating dNSP16 attenuation. Finally, we show that targeting NSP16 activity with a 2’-*O* methyltransferase inhibitor in combination with type I interferon offers a novel avenue for antiviral development.

## Introduction

Since its emergence late in 2019, severe acute respiratory syndrome coronavirus 2 (SARS-CoV-2) has caused major damage to the global populace through mortality (*1*), morbidity (*2*), and social and economic disruption (*3*). While the pandemic may be seen as shifting to endemicity, the continued threat of epidemic waves remains due to waning immunity and/or the emergence of new SARS-CoV-2 variants of concern (*4*). Moreover, future outbreaks caused by CoVs seem possible considering previous epidemics this century caused by SARS-CoV and Middle East respiratory syndrome (MERS)-CoV (*5*). Therefore, there is a need to expand our understanding of SARS-CoV-2 and identify additional avenues for treatment.

CoVs encode an array of viral effectors that subvert host immunity to allow for successful replication and pathogenesis (*6, 7*). However, variations in function and effect across the CoV family indicate a need to functionally test these effectors in viral replication and pathogenesis studies. CoV nonstructural protein (NSP16), a ribonucleoside 2’-*O* methyltransferase (MTase), catalyzes the transfer of a methyl group to the viral RNA cap structure (*8, 9*). This modification to the viral RNA cap is thought to prevent recognition by the host RNA sensor MDA5 and effectors in the interferon-induced protein with tetratricopeptide repeats (IFIT) family (*10, 11*). Reliance on 2’-*O* methylation has been observed in a broad range of virus families that either encode their own 2’-*O* MTases (*12*), rely on a host 2’-*O* MTase (*13*), or simply “snatch” host mRNA caps to incorporate into their own viral RNA (*14*). Disrupting the ability of these viruses to mimic host RNA cap structure results in a range of attenuation phenotypes (*10, 13, 15, 16*).

In this work, we confirmed the importance of SARS-CoV-2 NSP16 to viral infection and pathogenesis. Building from previous studies on CoV 2’-*O* MTases, we disrupted via mutagenesis a conserved lysine-aspartic acid-lysine-glutamate (KDKE) catalytic tetrad necessary for NSP16 MTase function (*11, 17*). We found the NSP16 MTase mutant (dNSP16) was attenuated *in vitro* in the context of type I interferon (IFN-I) activity. Additionally, we observed reduced disease and viral loads for dNSP16 in the hamster model. Importantly, we showed that the IFN-stimulated genes (ISGs) IFIT1 and IFIT3 mediate dNSP16 attenuation. Finally, targeting NSP16 activity with the MTase inhibitor sinefungin increased the sensitivity of wild-type (WT) SARS-CoV-2 to IFN-I treatment. Together, these findings demonstrate a key role for NSP16 in SARS-CoV-2 immune evasion and potentially identify CoV 2’-*O* MTase function as a target for novel therapeutic approaches (*18*).

## Results

### dNSP16 has no replication defect

To investigate the contribution of NSP16 to SARS-CoV-2, we constructed dNSP16 using our infectious clone of SARS-CoV-2 as previously described (*19, 20*). Briefly, we generated a 2-base pair substitution, converting aspartic acid to alanine (D130A) in the conserved KDKE motif (**Fig. 1a, b**). This mutation is predicted to ablate MTase function (*17*) and prior CoV studies have confirmed the importance of this residue to CoV replication and pathogenesis (*21-23*). We also attempted to construct an NSP16 deletion-virus by engineering an in-frame stop codon at the first amino acid position, but this deletion mutant failed to replicate. In IFN-deficient Vero E6 cells, dNSP16 displayed replication kinetics (**Fig. 1c**) and plaque sizes similar to those of WT (**Fig. 1d**). Together, these results suggest no significant impact on viral replicative capacity with the loss of NSP16 catalytic activity. Importantly, the D130A mutation was found to be stable in our rescued dNSP16 stock by Sanger sequencing and we confirmed no common spike mutations in the region adjacent to the furin cleavage site that have been previously reported for virus stocks amplified on Vero E6 cells (*24, 25*) (**Fig. S1**).

**Figure 1.**
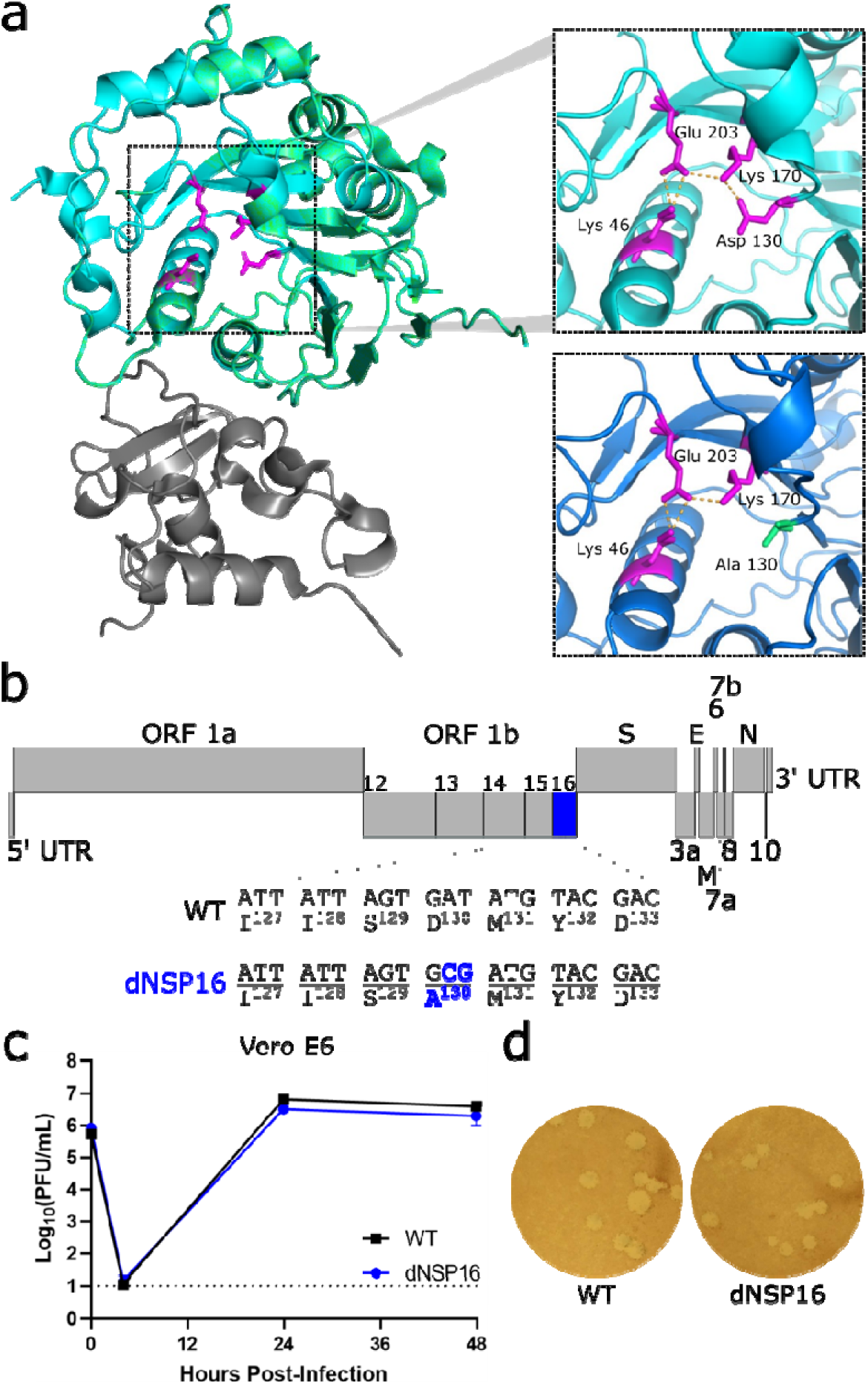
dNSP16 has no replication defect. (a) SARS-CoV-2 NSP16 (green) in complex with scaffold NSP10 (gray). The upper inset shows the KDKE catalytic tetrad (in magenta, with amino acids labeled) with polar contacts shown by orange dashed lines. The right panel shows mutation of the KDKE motif to KAKE (D130A). The structural modeling demonstrates a loss of a hydrogen bond between K170 and A130. Structures based on Protein Data Bank ID: 6W4H with homology model made using Swiss-Model (*18*). (b) Schematic of the SARS-CoV-2 genome, drawn to scale, with NSP16 highlighted in blue and the engineered two-base change indicated, resulting in coding change D130A. (c) Replication of WT (black) and dNSP16 (blue) in Vero E6 cells, multiplicity of infection = 0.01; *n* = 3. Means are plotted with error bars denoting standard deviation. Dotted line represents limit of detection. PFU = plaque-forming units. (d) Plaque morphology of the WT and dNSP16 viruses on Vero E6 cells.

### dNSP16 is attenuated in human respiratory cells

While the dNSP16 mutant had no replicative attenuation in Vero E6 cells, phenotypes in these cells are often not representative of relevant cells such as human respiratory cells (*25-27*). Therefore, we next evaluated dNSP16 in Calu-3 2B4 cells, a human lung carcinoma cell line. Compared to WT SARS-CoV-2, we observed significant attenuation of dNSP16 in Calu-3 2B4 cells (**Fig. 2a**). At both 24 and 48 hours post-infection (HPI), WT SARS-CoV-2 displayed robust replication whereas a 2.5 log_10_ decrease in replication was observed for dNSP16 at both time points. These results are consistent with similar findings for both SARS-CoV and MERS-CoV 2’-*O* MTase mutants (*15, 28*). Together, the results confirm the requirement of NSP16 for successful SARS-CoV-2 infection of human respiratory cells.

**Figure 2.**
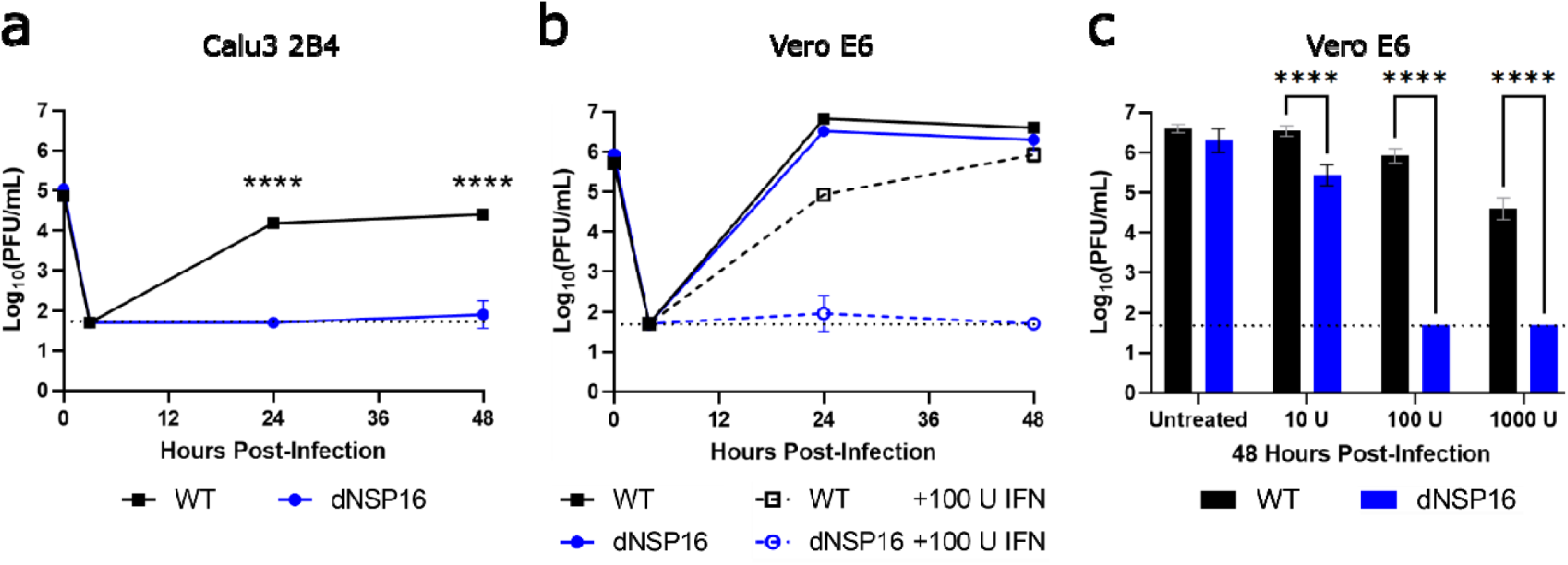
dNSP16 is attenuated in human respiratory cells and is more sensitive to type I interferon (IFN-I) pre-treatment. (a) Replication of WT (black) and dNSP16 (blue) in Calu-3 2B4 cells, MOI = 0.01. \*\*\*\**p*<0.001: results of two-way ANOVA with Tukey’s multiple comparison test (α = 0.05). (b) Replication of WT (black) and dNSP16 (blue) in Vero E6 cells without IFN-I (solid lines, data as in Fig. 1c), or with 100 U IFN pre-treatment a day prior to infection (dashed lines), multiplicity of infection = 0.01. (c) Comparison of the viral titers at 48 hours post-infection from panel (b), with additional treatment levels of IFN-I indicated. \*\*\*\**p*<0.001: results of two-way ANOVA with Tukey’s multiple comparison test (α = 0.05). Means are plotted with error bars denoting standard deviation. For all panels, *n* = 3 for all data points. Dotted lines represent limits of detection. PFU = plaque-forming units.

### dNSP16 is more sensitive to type I IFN pre-treatment

A major distinction between Vero E6 and Calu-3 2B4 cells is their capacity to induce a type I interferon (IFN-I) response; while Calu-3 2B4 cells are IFN-I competent, Vero E6 cells do not induce IFN-I, but do respond when treated exogenously. Therefore, we investigated the effects of IFN-I on the replication of dNSP16 relative to WT. Pre-treating Vero E6 cells with 100 U of IFN-I, we noted a modest, but significant decrease in WT infection compared to untreated cells (**Fig. 2b**). In contrast, Vero E6 cells pre-treated with IFN-I resulted in 3.0 log_10_ and 4.2 log_10_ decreases in dNSP16 titer at 24 and 48 HPI, respectively. We also observed a dose-dependent decrease in titer with respect to IFN-I pre-treatment for both dNSP16 and WT; however, the effect on dNSP16 was more pronounced, especially at higher IFN-I concentrations (**Fig. 2c**). Overall, the results indicate that dNSP16 is more sensitive to IFN-I compared to WT SARS-CoV-2.

### dNSP16 is attenuated *in vivo*

We next asked whether the attenuation of dNSP16 we observed *in vitro* would manifest *in vivo*. We challenged Syrian (golden) hamsters, a model for SARS-CoV-2 infection studies (*29*), intranasally (i.n.) with 10^4^ plaque-forming units (PFU) of dNSP16, WT, or a mock-infection control (**Fig. 3a**). While both dNSP16- and WT-infected hamsters showed weight loss relative to the mock-infected control hamsters, the dNSP16-infected hamsters showed reduced weight loss compared to WT-infected hamsters (**Fig. 3b**). Moreover, the dNSP16-infected hamsters did not show signs of disease, and only the WT-infected hamsters displayed ruffled fur at 5 and 6 days post-infection (DPI)(**Fig. 3c**). Lung histopathologic findings were more severe for WT-infected hamsters compared to dNSP16-infected hamsters at both 4 DPI and 7 DPI (**Fig. 3d**). Both groups developed interstitial pneumonia, bronchiolitis, peribronchiolitis, perivasculitis, and perivascular edema. WT-infected hamsters experienced a greater degree of subendothelial edema and hemorrhage. Together, these results indicate that dNSP16 results in reduced disease in the hamster model of SARS-CoV-2 infection.

**Figure 3.**
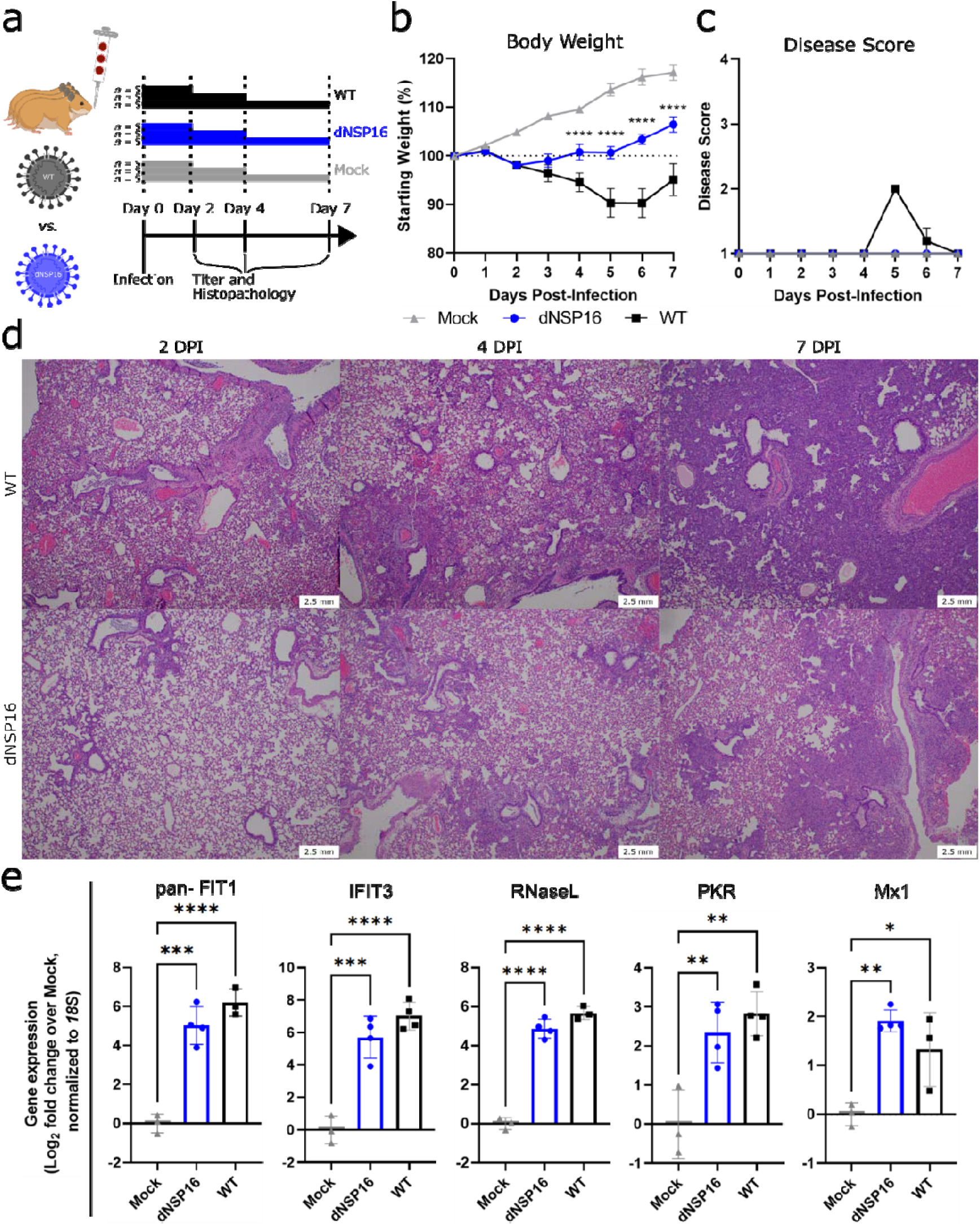
dNSP16 is attenuated *in vivo*. (a) Overview of experimental plan for hamster infections. 100 μL inoculum of PBS (mock) or either dNSP16 (10^4^ plaque-forming units) or WT (10^4^ plaque-forming units) was given intranasally to 4- to 5-week-old Syrian hamsters. At 2, 4, and 7 days post-infection (DPI), 5 animals from each infection group were sacrificed for organ collection. Some graphics generated from BioRender. (b) Percent starting weights and (c) disease scores for mock-, dNSP16-, or WT-infected hamsters. \*\*\*\**p*<0.001: results of a mixed-effects model (restricted maximum likelihood) with Tukey’s multiple comparison test (α = 0.05) performed between WT- and dNSP16-infected hamsters at the indicated DPI. Means are plotted with error bars denoting standard error of the mean. (d) Hematoxylin and eosin staining of representative 5 μm-thick sections taken from left lung lobes. (e) Fold change (log_2_) of expression of the indicated immune genes from right middle lung lobes isolated from hamsters infected with the indicated virus (or mock), 2 DPI. For each panel, fold changes from dNSP16 or WT samples are measured relative to mock samples. Values from individual hamsters are plotted (symbols) as well as means (bars). Error bars denote standard deviation. All samples were normalized to 18S expression, used as a reference. \**p*<0.05, \*\**p*<0.01, \*\*\**p*<0.005, \*\*\*\**p*<0.001: results of one-way ANOVA with Tukey’s multiple comparison test (α = 0.05).

To explore why disease phenotype differed in dNSP16-infected hamsters, we first evaluated changes in the host immune response following infection with dNSP16. Examining RNA from hamster lungs collected at 2 DPI, we observed that both WT- and dNSP16-infected samples had increased expression of ISGs (IFIT1, IFIT3, RNase L, PKR, and Mx1) as well as other immune genes (IFNγ, IL-1β, IL-10) (**Fig. S2**) relative to mock. However, no differences in gene expression were observed between WT- and dNSP16-infected hamsters; these data correspond to previous findings with a SARS-CoV 2’-*O* MTase mutant (*15*). Our results suggest the loss of NSP16 activity may not drive increased immune gene expression, but rather sensitize dNSP16 to immune gene activity otherwise ineffective against WT SARS-CoV-2.

### dNSP16 replication is reduced *in vivo*

We next evaluated viral load in dNSP16-infected versus WT-infected hamsters. Examining replication in the lung, we observed similar viral loads at 2 DPI between dNSP16- and WT-infected hamsters (**Fig 4a**); however, by 4 DPI, dNSP16 titer was significantly reduced. This delayed attenuation in the lung corresponds to previous reports for both SARS-CoV and MERS-CoV in mice (*28, 30*). However, nasal wash titers at both 2 and 4 DPI were lower for dNSP16-compared to WT-infected hamsters (**Fig. 4b**). These nasal wash titer data suggest attenuation of dNSP16 occurs in the upper airway at an earlier time compared to lung and suggest different tissue-mediated immune responses between the upper and lower respiratory tract. Notably, while viral titers in the lung were equivalent at 2 DPI, nucleocapsid-specific staining of lung tissue showed more pervasive staining for WT-compared to dNSP16-infected tissues (**Fig. 4c**). This trend was exacerbated at 4 DPI and corresponded to the difference in titer observed between dNSP16- and WT-infected hamsters (**Fig. 4a**). Consistent with differences in fitness *in vivo*, targeted Sanger sequencing of viral RNA from the lungs at 4 DPI showed no signs of reversion in the dNSP16-infected hamsters (**Fig. S3**). Together, these results indicate that dNSP16 causes reduced disease and exhibits decreased viral replication *in vivo* despite inducing an immune response similar to that of WT SARS-CoV-2.

**Figure 4.**
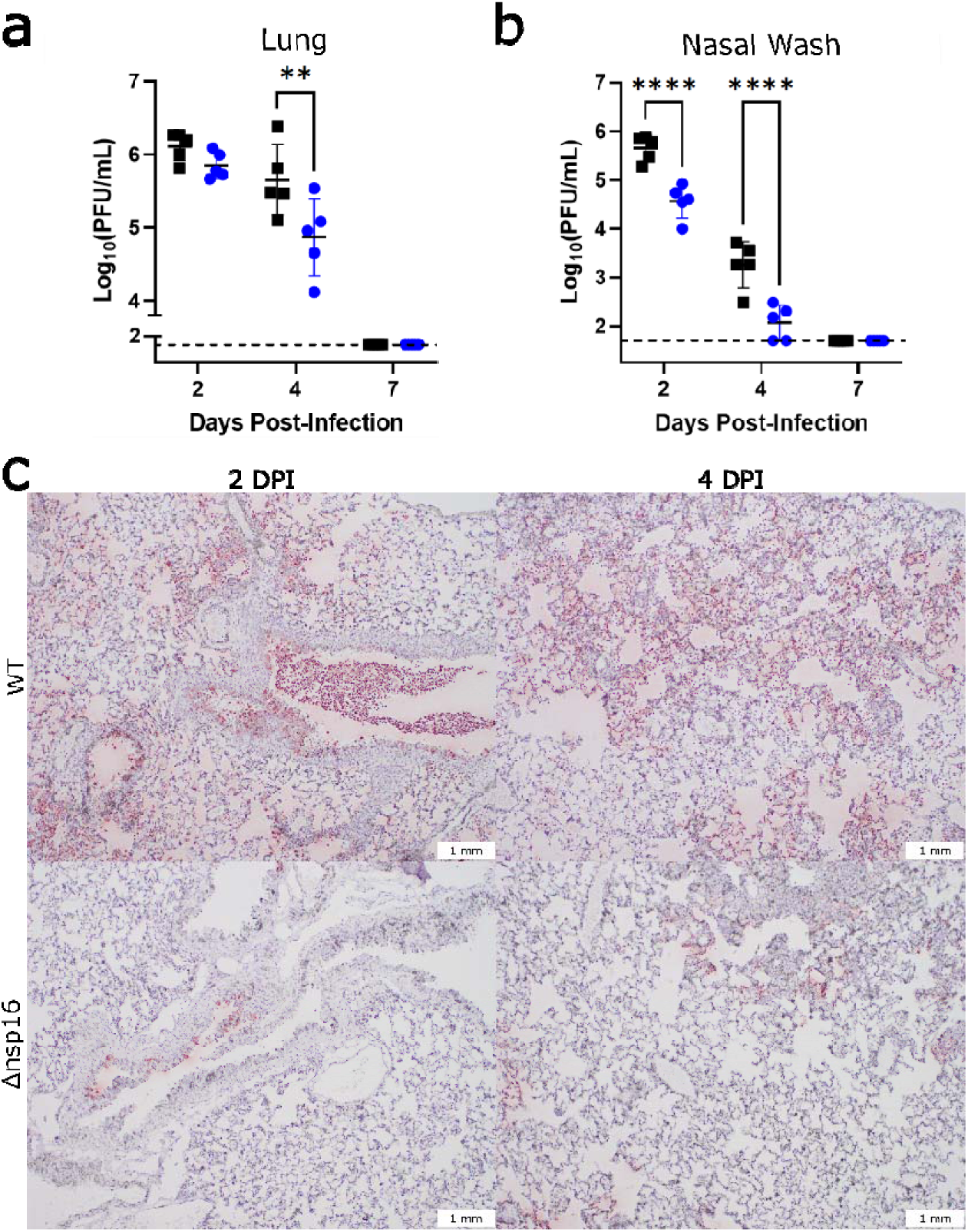
dNSP16 replication is reduced *in vivo*. (a, b) Comparison of viral titers from (a) right cranial lung lobes or (b) nasal washes from WT- (black) or dNSP16-infected (blue) hamsters sacrificed at the indicated day. \*\**p*<0.01, \*\*\*\**p*<0.001: results of two-way ANOVA with Tukey’s multiple comparison test (α = 0.05). Values from individual hamsters are plotted (symbols) as well as means (black bars). Error bars denote standard deviation. Dotted lines represent limits of detection. PFU = plaque-forming units. (c) SARS-CoV-2 nucleocapsid staining (brown) of representative 5 μm-thick sections taken from left lung lobes.

### Knockdown of IFIT genes partially reverses attenuation of dNSP16

Based on increased sensitivity to IFN-I, attenuation of dNSP16 is likely mediated by sensitivity to certain ISG effectors. Therefore, we focused on several ISGs known to target foreign nucleic acids including the IFIT family (*31*), PKR (*32*), and OAS1 (*33*). We transfected Vero E6 cells with target or control siRNAs, treated them with IFN-I, and then infected with either WT SARS-CoV-2 or dNSP16. Whereas control siRNA treatment resulted in undetectable viral titers for dNSP16 at 48 HPI, consistent with the attenuating effect of IFN-I (**Fig. 2b, c**), we observed a significant restoration of viral titers with anti-IFIT1 siRNA treatment (**Fig. 5a**). Similarly, siRNA-induced knockdown of IFIT3, shown to stabilize IFIT1 and enhance its cap-binding function (*34*), resulted in a restoration of dNSP16 titers comparable to those observed with anti-IFIT1 siRNA. However, the combination of IFIT1 and IFIT3 knockdown had no additive impact in these studies. Notably, neither anti-PKR nor anti-OAS1 siRNA treatment significantly affected dNSP16 replication relative to control siRNA despite confirming knockdown for all targets (**Fig. S4**). Together, the results suggest that both IFIT1 and IFIT3 play critical roles in the attenuation of dNSP16.

**Figure 5.**
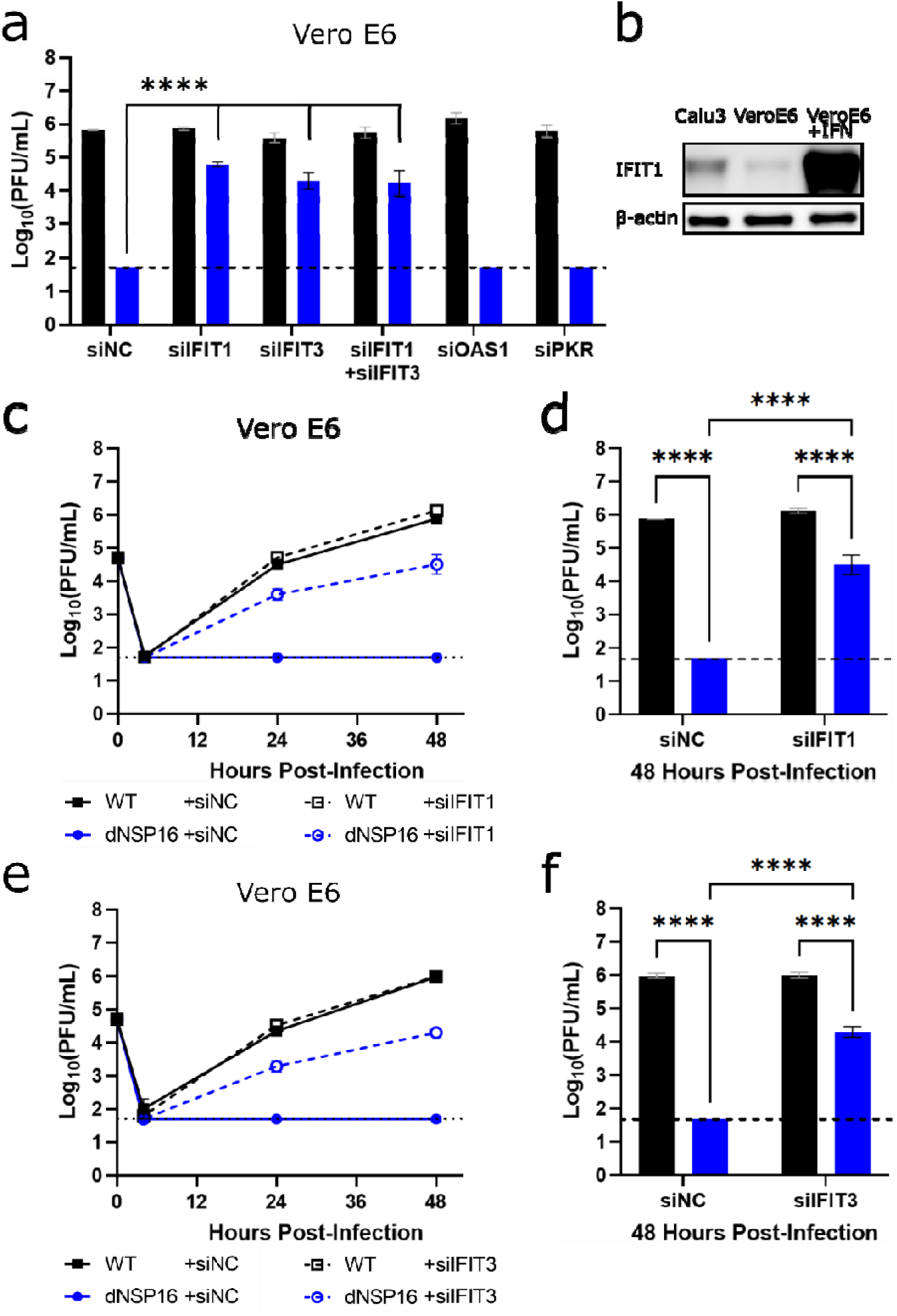
Knockdown of IFIT genes partially reverses attenuation of dNSP16. (a) Replication of WT (black) and dNSP16 (blue) in the context of siRNA treatment. 1.25 × 10^5^ Vero E6 cells/well were reverse transfected with 2 pmol total of the indicated siRNA construct(s) 2 days prior to infection and also pre-treated with 100 U IFN-I a day prior to infection, multiplicity of infection (MOI) = 0.01. Data shown at 48 hours post-infection (HPI). Statistical comparisons on graph are with respect to siRNA control treatment (“siNC”). (b) Baseline IFIT1 protein expression in Calu-3 2B4 and Vero E6 cells, or Vero E6 cells 1 day post-stimulation with IFN-I. (c) Viral replication kinetics for WT (black) or dNSP16 (blue) following treatment with anti-IFIT1 (dashed) or control siRNA (solid). 1.25 × 10^5^ Vero E6 cells were reverse transfected with 1 pmol of the indicated siRNA construct 2 days prior to infection and also pre-treated with 100 U IFN-I a day prior to infection, MOI = 0.01. (d) Comparison of the viral titers at 48 HPI from panel (c), black = WT, blue = dNSP16. (e) Viral replication kinetics for WT (black) or dNSP16 (blue) following treatment with anti-IFIT3 (dashed) or control siRNA (solid). 1.25 × 10^5^ Vero E6 cells/well were transfected with 1 pmol of the indicated siRNA construct 2 days prior to infection and also pre-treated with 100 U IFN-I a day prior to infection, MOI = 0.01. (f) Comparison of the viral titers at 48 HPI from panel (e), black = WT, blue = dNSP16. For panels (a), (d) and (f), \*\*\*\**p*<0.001: results of two-way ANOVA with Tukey’s multiple comparison test (α = 0.05). Means are plotted with error bars denoting standard deviation. For all panels, *n* = 3 biological replicates for all data points. Dotted lines represent limits of detection. PFU = plaque-forming units.

IFIT family members have previously been shown to recognize non-host mRNA cap structures (*35*). Based on the initial siRNA screen (**Fig. 5a**), we next evaluated if the differences in viral attenuation we noted between dNSP16 and WT SARS-CoV-2 may be due to the presence of baseline IFIT1 expression in the cells we tested. We subsequently observed that Calu-3 2B4 cells expressed IFIT1 protein at baseline, whereas expression of IFIT1 in Vero E6 cells was low (**Fig. 5b**). However, upon stimulation of Vero E6 cells with IFN-I, we observed a robust induction of IFIT1 that may account for the dNSP16 attenuation we noted (**Fig. 2c**). We further examined the replication kinetics of dNSP16 in the context of IFIT1 knockdown (**Fig. 5c**). Whereas treatment with 100 U of IFN-I and control siRNA resulted in undetectable viral titers for dNSP16 at all time points tested, we observed partial restoration of viral titers for dNSP16 in the context of anti-IFIT1 siRNA treatment at both 24 and 48 HPI (**Fig 5c, d**). While the role of IFIT1 has previously been noted for CoV 2’-*O* MTases (*15, 28*), IFIT3 has only recently been shown to enhance IFIT1’s RNA-binding ability in human cells (*34*). Similar to IFIT1 knockdown, IFIT3 knockdown restored replication of dNSP16 at both 24 and 48 HPI (**Fig. 5e, f**). Since IFIT1 and IFIT3 share sequence homology, we also confirmed that both our anti-IFIT1 and anti-IFIT3 siRNA constructs were specific to their respective targets (**Fig. S5**). Coupled with the fact that combined anti-IFIT1/anti-IFIT3 siRNA treatment had no additive effect (**Fig. 5a**), these results suggest both human IFIT1 and IFIT3 are necessary for attenuation of SARS-CoV-2 dNSP16.

### Targeting the NSP16 active site for antiviral treatment

Having established the critical role for NSP16 in helping SARS-CoV-2 evade IFIT function, we next explored whether NSP16 activity could be targeted for therapeutic treatment. Using sinefungin, an *S*-adenosyl-L-methionine (SAM) analogue and inhibitor of SAM-dependent MTases (*36*), we attempted to disrupt NSP16 MTase activity and reduce replication of WT SARS-CoV-2. Previous modeling studies demonstrated that sinefungin binds in the active site of NSP16, interacting with the D130 residue we mutated in dNSP16 (**Fig. 6a**) (*37*). We tested a range of sinefungin concentrations on WT SARS-CoV-2 replication in Vero E6 cells. We observed a dose-dependent decrease in SARS-CoV-2 replication, with 5 mM and 10 mM concentrations reducing replication by 1.6 log_10_ and 3.1 log_10_, respectively (**Fig. 6b**, solid bars). Together, the results suggest that sinefungin treatment acts on viral MTase function to attenuate viral replication.

**Figure 6.**
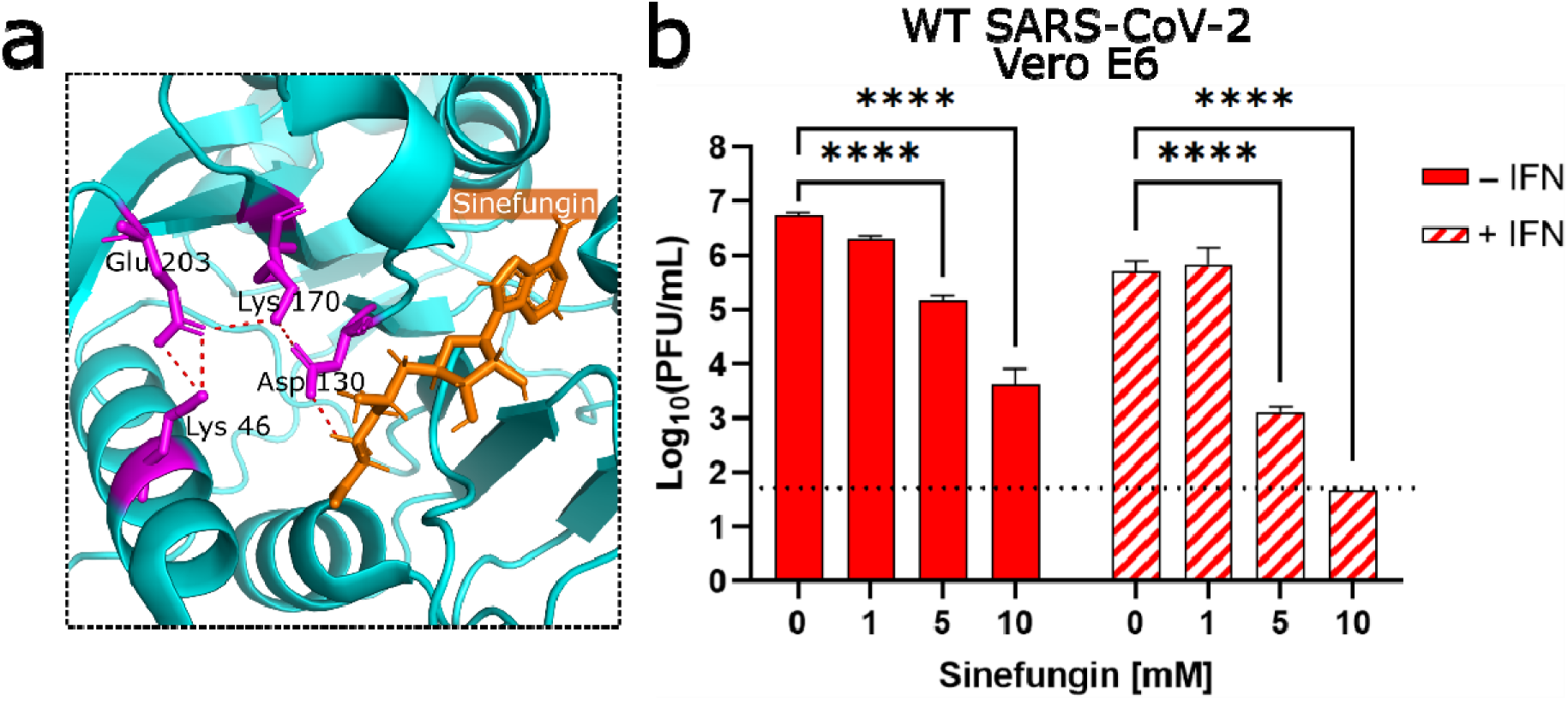
Targeting the NSP16 active site for antiviral treatment. (a) Detail of structure of NSP16 in complex with sinefungin, from Protein Data Bank ID: 6YZ1 (*37*). The residues of the catalytic core are colored in magenta, sinefungin is colored in orange, and polar contacts are shown by orange dashed lines. (b) Dose-dependent effect of sinefungin on WT SARS-CoV-2 replication. 5 × 10^4^ Vero E6 cells/well were seeded in 24-well format one day before infection and also pre-treated with 100 U IFN-I 8 hours later.The day of infection (multiplicity of infection = 0.01), sinefungin was given at the indicated concentration 1 hour after infection (in cell culture media). Data shown at 48 HPI. \*\*\*\**p*<0.001: results of two-way ANOVA with Tukey’s multiple comparison test (α = 0.05).Means are plotted with error bars denoting standard deviation. *n* = 3 biological replicates for all data points. PFU = plaque-forming units.

While IFN-I treatments (both IFNα and IFNβ) have significant impacts in randomized clinical trials (*38*), our earlier data suggests that disruption of NSP16 activity will sensitize SARS-CoV-2 to IFN-I-induced effectors like IFIT1 and IFIT3. Therefore, we tested the additive impact of sinefungin and IFN-I pre-treatment used in combination (**Fig. 6b**, striped bars). Our results indicated that this combination treatment drove attenuation of WT-SARS-CoV-2 to the replication levels observed with dNSP16 (**Fig. 2b**). IFN-I treatment alone resulted in a modest, but significant reduction in titer (1.0 log_10_), consistent with earlier data (**Fig. 2b**). However, the addition of sinefungin to IFN-I pre-treatment resulted in a sinefungin dose-dependent reduction in titer beyond that induced by IFN-I alone. Notably, treatment with 10 mM sinefungin with IFN-I resulted in SARS-CoV-2 titers near the limit of detection, a 5.0 log_10_ drop in titer compared to mock-treated cells. Together, the results argue that combination approaches which target NSP16 to sensitize SARS-CoV-2 to IFN-I responses may offer a novel approach for therapeutic CoV treatments.

## Discussion

In this study, we engineered NSP16-mutant SARS-CoV-2 with an amino acid change at a conserved catalytic residue, D130A. The mutant, dNSP16, replicated similarly to WT SARS-CoV-2 in the IFN-I-deficient cell line Vero E6, but was attenuated in human respiratory cells. Moreover, dNSP16 showed greater sensitivity to pre-treatment with exogenous IFN-I compared to WT. *In vivo*, dNSP16 was attenuated compared to WT, as evidenced by decreased weight loss, lack of clinical signs of disease, and reduced pathologic changes in the hamster lung. Attenuated disease corresponded to lower viral titers in the nasal wash and lung, as well as reduced viral antigen staining in the lung. Mechanistically, the attenuation of dNSP16 is mediated by IFIT1 and IFIT3, with knockdown of either gene restoring viral replication in the context of IFN-I pre-treatment. Lastly, we found that sinefungin, an *S*-adenosyl-L-methionine (SAM) analogue that targets NSP16 activity, reduced WT SARS-CoV-2 replication. In addition, the effect of sinefungin on reducing viral replication was enhanced when combined with IFN-I pre-treatment, likely as a result of decreased NSP16 MTase function and a corresponding increase in recognition by IFIT proteins. Together, our work highlights the critical role of NSP16 in neutralizing the antiviral effects of IFIT1/IFIT3 against WT SARS-CoV-2.

Ablating NSP16 MTase activity does not result in loss of the replicative capacity of dNSP16 compared to WT. Yet, our inability to rescue an NSP16-deletion virus with an inserted stop codon suggests NSP16’s role may be more complex than its 2’-*O* MTase activity alone. Notably, replication attenuation of dNSP16 occurs in the context of a viable IFN-I response. These results are consistent with previous studies of 2’-*O* MTase mutants in CoVs including SARS-CoV (*15*) and MERS-CoV (*28*). Similarly, reduced disease and attenuation of viral replication at 4 DPI in the lung of dNSP16-infected hamsters is consistent with data from other 2’-*O* MTase CoV mutants in mouse models (*15, 28*). However, our viral titer data from nasal washes, a measure of viral fitness in the upper airway, indicate that dNSP16 attenuation occurs at the earlier 2 DPI time point. These results, not surveyed in the CoV mouse models, suggest the upper airway and the lung have distinct immune activation responses, leading to different kinetics for dNSP16 attenuation.

Our studies also confirm a role for the IFIT proteins in mediating attenuation of dNSP16. Previously, human IFIT1, an ISG, has been shown to sequester viral mRNA lacking 2’-*O* methylation (*39*) through a mechanism that involves direct recognition of the cap structure (*40*). In prior studies with CoVs, mouse Ifit1, a paralog to human IFIT1 (*41*), antagonized CoVs lacking 2’-*O* methylation (*10, 11*). Here, we demonstrate that while dNSP16 is attenuated by IFN-I pre-treatment in Vero E6 cells, knockdown of IFIT1 partially restores dNSP16 replication. In addition, rapid attenuation of dNSP16 in Calu-3 2B4 cells, compared to Vero E6 cells, may be due to higher baseline levels of IFIT1 in the former. We also found that knockdown of IFIT3 partially restored dNSP16 replication in the context of IFN-I pre-treatment. Recent studies have highlighted the importance of IFIT3 in stabilizing IFIT1 function and optimizing its recognition of RNA caps lacking 2’-*O* methylation (*34*). Notably, the combination of IFIT1 and IFIT3 knockdown we tested had no additive effect, suggesting that both together are required for restriction of dNSP16. Overall, these results indicate the importance of NSP16 in protecting CoVs from IFIT effector function.

Having established a critical role for NSP16 in evading IFIT activity, we evaluated the feasibility of targeting 2’-*O* methylation of CoVs therapeutically. Using sinefungin, a pan-inhibitor of SAM-dependent MTases, we observed a dose-dependent reduction in replication of WT SARS-CoV-2, indicating that targeting viral MTases activity can impair successful infection. Importantly, combined treatment with sinefungin and IFN-I had an additive effect, resulting in increased attenuation, likely due to both a loss of viral 2’-*O* methylation and increased recognition of unmethylated viral RNA by IFIT1/IFIT3. This approach is distinct from those of other CoV therapies targeting the viral polymerase (*42*) or the main protease (*43*) to arrest virus replication. Targeting NSP16 similarly disrupts a viral enzymatic process, yet here, an effector response is provided by the host via IFIT proteins. Importantly, while attenuation of dNSP16 is delayed in the hamster lung, early attenuation in the upper airway suggests more rapid or robust expression of IFIT proteins in the upper airway. This could, in turn, increase the efficacy of drugs targeting CoV 2’-*O* MTase activity in the upper airway, a possible strategy to decrease transmission and spread. With augmented upper airway replication as a feature of SARS-CoV-2 variants of concern (*44*), NSP16-targeting drugs may provide an effective countermeasure for the current and future CoV pandemics.

Overall, our results confirm the importance of NSP16 to SARS-CoV-2 infection and pathogenesis. A mutation that disrupts the NSP16 2’-*O* MTase catalytic site attenuates disease *in vivo* and demonstrates its importance in evading host innate immunity. In the absence of 2’-*O* MTase activity, SARS-CoV-2 is rendered susceptible to the effector responses of IFIT1 and IFIT3 in combination. Importantly, such dependence of SARS-CoV-2 on the 2’-*O* MTase function of NSP16 offers a novel target for future CoV antiviral drug development.

## Acknowledgments

We thank Eileen McAnarney for her prior contributions to the laboratory, which facilitated this work. We also thank Jordyn Walker for her help in ear-tagging our hamsters. Figures were created with BioRender.com and Inkscape.

## Author contributions

Conceptualization: CS, VDM

Formal analysis: CS, PACV, SS, KD, ALR, DHW

Funding acquisition: CS, ALR, SCW, VDM

Investigation: CS, DS, JAP, BK, SS, KSP

Methodology: CS, KL, MNV, BAJ, DS, JAP, BK, SS, REA, MDD, KSP, VDM

Project Administration: VDM

Supervision: ALR, SCW, MDD, KSP, VDM

## Materials and Methods

### Cells

Vero E6 cells (ATCC #CRL-1586) were cultured in high-glucose Dulbecco’s Modified Eagle Medium (DMEM, Gibco #11965–092) supplemented with 5% heat-inactivated fetal bovine serum (FBS, Cytiva #SH30071.03) and 1X Antibiotic-Antimycotic (Gibco #15240-062). VeroE6/TMPRSS2 (JCRB #1819) were cultured in low-glucose, pyruvate-containing DMEM (Gibco #11885-084) supplemented with 5% FBS and 1 mg/mL geneticin (Gibco #10131-035). Calu-3 2B4 (BEI Resources # NR-55340) were cultivated in high-glucose DMEM supplemented with 10% FBS, 1X Antibiotic-Antimycotic, and 1 mM sodium pyruvate (Sigma-Aldrich #S8636). Baby hamster kidney (BHK) cells were cultured in MEM α with GlutaMAX (Gibco # 32561-037) supplemented with 5% FBS and 1X Antibiotic-Antimycotic. For all propagation and experimentation, cells were kept at 37°C and 5% CO_2_ in a humidified incubator.

### Viruses

We performed PCR-based mutagenesis to engineer a 2-base pair (bp) mutation in codon 130 of the NSP16 gene encoded on a SARS-CoV-2 infectious clone (ic) reverse genetics system based on the prototype “USA/WA1/2020” strain (NCBI accession no.: MN985325), following our previously published method (*19, 20*). The engineered change was made to the second and third bp positions of NSP16 codon 130 (*GAT*➔*GCG*) on pUC57-CoV-2-F5, changing the encoded aspartic acid residue to an alanine. The initially rescued virus constituted a heterogenous population of sequences, therefore the initial stock was serially diluted and plated into wells containing Vero E6 cells to isolate single clones via plaque purification. Individual plaques were carefully scraped with a pipette tip and used to inoculate separate wells containing Vero E6 cells. Upon induction of CPE, culture supernatants were cleared of cellular debris and part of the liquid fraction processed for viral RNA purification and Sanger sequencing. Well supernatants associated with viral sequences that contained the desired NSP16 mutation were then used to infect TMPRSS2-expressing Vero E6 cells for an additional round of virus replication to generate higher viral titers; TMPRSS2-expressing cells were chosen to reduce the chance of mutation of the spike protein around the furin cleavage site (*24*). The supernatants from these cells were similarly processed as described above for confirmation of viral sequence via Sanger sequencing. Upon sequence verification, a supernatant-stock of icSARS-CoV-2 with the engineered NSP16 mutation (“dNSP16”) was selected for use in subsequent experiments. With the exception of the plaque purification step, wild-type icSARS-CoV-2 (“WT”) was produced in the same way as dNSP16.

### Viral replication kinetics

Cells were seeded in 24-well format. In experiments involving IFN-I pre-treatment, cells were treated 16 – 20 hours prior to infection with Universal Type I IFN (PBL Assay Science #11200-2), diluted in Dulbecco’s phosphate-buffered saline, without calcium chloride and magnesium (DPBS, Gibco #14190-144). After infection at a multiplicity of infection (MOI) of 0.01 and incubation for 45 minutes at 37°C with 5% CO_2_ and manual tilting every 15 minutes, cells were washed 3X with 500 μL DPBS and then given 500 μL of cell type-specific medium. Supernatants were collected within 1 hour of the indicated time point whereupon 150 μL of culture medium was removed and an equal volume of fresh medium was added back to the sample well. Supernatant samples were subsequently titered via plaque assay. All conditions were performed in triplicate, and all experiments were performed in an approved biosafety level 3 (BSL3) laboratory at the University of Texas Medical Branch at Galveston (UTMB).

### Plaque assay

One day before the assay, 6-well plates were seeded with 3 × 10^5^ Vero E6 cells/well. Under BSL3 conditions, samples of virus-containing supernatant were titrated in a 10-fold dilution series in DPBS, and 200 μL of each dilution of the series was transferred to confluent cells after culture medium was removed. Assay plates were incubated at 37°C with 5% CO_2_ for 45 minutes with manual tilting every 15 minutes. Afterwards, an overlay of 1X Modified Eagle Medium (Gibco #11935-046) containing 5% heat-inactivated FetalClone II (Cytiva #SH30066.03), 1X Antibiotic-Antimycotic, and 1% agarose (Lonza #50004) was applied to wells, and the plates were returned to the incubator for two days. Afterwards, a 1X dilution in DPBS of 10X neutral red stain (0.85% w/v NaCl, 0.5% w/v Fisher Scientific #N129-25) was applied to each well, and 2 – 5 hours later, plaque-forming units (PFU) were visualized using a lightbox and manually counted. The limit of detection was 50 PFU/mL, corresponding to 1 PFU in the well with the lowest dilution factor (1:50 total dilution).

### Animal studies

Four- to five-week-old male Syrian hamsters (*Mesocricetus auratus*), strain HsdHan:AURA, purchased from Envigo were infected intranasally (i.n.) with a 10^4^ PFU dose of either dNSP16 or WT in a 100 uL inoculum volume, or DPBS for mock-infected animals. Hamsters were randomly assigned to different treatment groups. Animal weights and clinical signs were recorded daily for up to 7 days post-infection (DPI). Disease scores were as follows: 1 (healthy), 2 (ruffled fur), 3 (hunched posture, orbital tightening, lethargy), 4 (moribund). At 2, 4, and 7 DPI, nasal washes from 5 animals from each experimental group were collected and the animals subsequently sacrificed, with right cranial, right middle, and left lung lobes from each animal collected in either DPBS, RNAlater (Invitrogen #AM7021), or 10% phosphate-buffered formalin (Fisher #SF100) for subsequent analyses of viral titer, gene expression and viral sequence, or histopathology, respectively. For measurement of viral titer, collected lung lobes were homogenized at 6000 rpm for 60 seconds using a Roche MagNA Lyser instrument and then titered via plaque assay. For analysis of gene expression and viral sequence, lung lobes stored in RNAlater were transferred to TRIzol (Invitrogen #15596018) and homogenized 5 times at 6500 rpm for 30 seconds, with cooling on a −20°C-chilled rack for 1 minute between homogenization steps. The homogenates were then processed for RNA purification as described below. For histopathological analysis, lung lobes were incubated with 10% phosphate-buffered formalin for 7 days at 4°C to allow for deactivation and buffer exchange before processing. All animal handling was performed at animal biosafety level 3 (ABSL3) conditions and in accordance with guidelines set by the Institutional Animal Care and Use Committee (IACUC) of the University of Texas Medical Branch.

### Histology

For visualization of histopathology, sections of paraffin-embedded formalin-fixed tissue were stained with hematoxylin and eosin on a SAKURA VIP 6 tissue processor at the University of Texas Medical Branch Surgical Pathology Laboratory. For visualization of viral antigen, tissue sections were deparaffinized and stained with a SARS-CoV-2 N-specific rabbit monoclonal antibody (Sino Biological #40143-R001) at a dilution of 1:30,000 followed by an anti-rabbit HRP-linked secondary (Cell Signaling #7074). Signal was developed with ImmPact NovaRED peroxidase kit (Vector Labs # SK-4805).

### RNA purification

RNA from cell supernatants, cell lysates, or homogenized lung tissue was extracted in TRIzol LS (Invitrogen #10296010) for cell supernatants only or TRIzol, followed by purification using Direct-zol RNA Miniprep Plus (Zymo Research #R2072) and reverse transcription using iScript cDNA synthesis kit (Bio-Rad #1708891).

### Sanger sequencing

Phusion High-Fidelity PCR Master Mix with HF Buffer (New England BioLabs #M0530) was used to amplify cDNA around the region of interest. 45 amplification cycles were used; otherwise the manufacturer’s protocol was followed. To amplify the region encoding NSP16, forward primer 5’-AACAGATGCGCAAACAGG and reverse primer 5’-TGCAGGGGGTAATTGAGTTC were used. To amplify the region of spike in the vicinity of the furin cleavage site, forward primer 5’-AGGCACAGGTGTTCTTAC and reverse primer 5’-TGAAGGCTTTGAAGTCTGCC were used. Amplicons were verified by gel electrophoresis, purified using QIAquick PCR Purification Kit (QIAGEN #28106), and sent to Genewiz (South Plainfield, NJ) for Sanger sequencing.

### Gene expression via quantitative PCR (qPCR)

qPCR was performed on cDNA using Luna (New England BioLabs #M3003) according to the manufacturer’s instructions. Fluorescent readings were made on a Bio-Rad CFX Connect instrument using Bio-Rad CFX Maestro 1.1 software (version 4.1.2433.1219). Relative gene expression was calculated manually using the ΔΔCt method: For each cDNA sample, the threshold cycle (Ct) of the gene of interest was first normalized against the Ct of the indicated reference gene. Then, the fold change in normalized expression for the gene of interest in each sample was calculated relative to normalized expression of the gene of interest in the control sample. The primers used for amplifying hamster targets were: 18S (forward: 5’ – GTAACCCGTTGAACCCCATT; reverse: 5’ – GTAACCCGTTGAACCCCATT), pan-IFIT1 (predicted to amplify NCBI accession nos.: XM_021224958, XM_040745240, and XM_013110344, forward: 5’ – TGCAGAGCTTGAAAGAAGCA; reverse: 5’ – CCTTCCTCACAGTCCACCTC), IFIT3 (forward: 5’ – CCTGGAGTGCTTAAGGCAAG; reverse: 5’ – TGCCTCACCTTGTCCACATA), RNase L (forward: 5’ – CCAGAGGGTAAAAACGTGGA; reverse: 5’ – TGCACCAAACCTGTGTGTTT), PKR (forward: 5’ – AAGTGCGTGAAGTAAAGGCG; reverse: 5’ – ATCCATTGCTCCAGAGTCCC), Mx1 (forward: 5’ – CTTCAAGGAGCACCCACACT; reverse: 5’ CTTGCCCTCTGGTGACTCTC), IFNγ (forward: 5’ – GGCCATCCAGAGGAGCATAG; reverse: 5’ – TTTCTCCATGCTGCTGTTGAA), IL-1β (forward: 5’ – GGCTGATGCTCCCATTCG; reverse: 5’ – CACGAGGCATTTCTGTTGTTCA), IL-10 (forward: 5’ – GTTGCCAAACCTTATCAGAAATGA; reverse: 5’ – TTCTGGCCCGTGGTTCTCT). The primers used for amplifying targets in Vero E6 cells were: β-actin (5’ – GGCATCCTCACCCTGAAGTA, reverse: 5’ – GGGGTGTTGAAGGTCTCAAA), IFIT1 (forward: 5’ – ACACCTGAAAGGCCAGAATG; reverse: 5’ – GCTTCTTGCAAATGTTCTCC), IFIT3 (forward: 5’ AGGAAGGGTGGACACAACTG; reverse: 5’ – TGGCCTGTTTCAAAACATCA), OAS1 (forward: 5’ – GATCTCAGAAATACCCCAGCCA; reverse: 5’ – AGCTACCTCGGAAGCACCTT), PKR (forward: 5’ – ACGCTTTGGGGCTAATTCTT; reverse: 5’ – TTCTCTGGGCTTTTCTTCCA). All primers were purchased as single-stranded DNA oligomers purified with standard desalting (Integrated DNA Technologies, Coralville, Iowa).

*DsiRNA experiments*. The following dicer-substrate short interfering RNAs (DsiRNAs)(Integrated DNA Technologies) were utilized: anti-IFIT1 (sense: 5’ – rGrCrUrUrGrArGrCrCrUrCrCrUrUrGrGrGrUrUrCrGrUrCTA; antisense: 5’ – rUrArGrArCrGrArArCrCrCrArArGrGrArGrGrCrUrCrArArGrCrUrU), anti-IFIT3 (sense: 5’ – rArGrCrUrGrArGrUrCrCrUrGrArUrArArCrCrArArUrArCGT; antisense: 5’ – rArCrGrUrArUrUrGrGrUrUrArUrCrArGrGrArCrUrCrArGrCrUrCrA), anti-OAS1 (sense: 5’ – rCrGrGrUrCrUrUrGrGrArArUrUrArGrUrCrArUrArArArCTA; antisense: 5’ – rUrArGrUrUrUrArUrGrArCrUrArArUrUrCrCrArArGrArCrCrGrUrC), anti-PKR (sense: 5’ – rGrUrArUrUrGrGrUrArCrArGrGrUrUrCrUrArCrUrArArACA; antisense: 5’ – rUrGrUrUrUrArGrUrArGrArArCrCrUrGrUrArCrCrArArUrArCrUrA), and negative control DsiRNA (Integrated DNA Technologies #51-01-14-03). For DsiRNA experiments, 1.25 × 10^5^ Vero E6 cells/well were reverse transfected in 24-well plate format with 1 – 2 pmol/well DsiRNA as indicated, 2 days prior to infection. 16 – 20 hours prior to infection cells were treated with 100 U of DPBS-diluted Universal Type I IFN (PBL Assay Science #11200-2). Infections proceeded as described in the section “viral replication kinetics” above.

### Protein expression via western blot

Cell lysates were harvested with 2X Laemmli SDS-PAGE sample buffer (Bio-Rad #1610737) containing a final concentration of 5% β-mercaptoethanol (Bio-Rad #1610710). Cell lysates were then denatured at 95°C for 10 min. The lysates were then loaded onto a Mini-PROTEAN TGX gel (Bio-Rad #4561096) and electrophoresed, followed by transfer to a polyvinylidene difluoride membrane (Bio-Rad #1620177). The membrane was then blocked in 5% nonfat dry milk dissolved in Tris-buffered saline with 0.1% Tween-20 (TBS-T) for 1 hour, followed by a short TBS-T wash. Overnight incubation with primary antibody, either rabbit anti-hIFIT1 (Cell Signaling Technology #14769) or rabbit anti-β-actin (Cell Signaling Technology #4970) was then performed. After, the membrane was washed 3 times with TBS-T and incubated with horseradish peroxidase-conjugated secondary antibody (Cell Signaling Technology #7074) for 1 hour. Finally, the membrane was washed 3 times with TBS-T, incubated with Clarity Western ECL Substrate (Bio-Rad #1705060), and imaged with a Bio-Rad ChemiDoc Imaging System running Bio-Rad Image Lab Touch software (version 2.4.0.03).

### Statistics

All statistics were performed in GraphPad Prism 9 (version 9.0.2), with details given in figure legends. Two-way ANOVA was performed on log_10_-transformed viral titers, with Tukey’s multiple comparison test (α = 0.05) to infer significant differences. For qPCR data, one-way ANOVA was performed on log_2_-transformed ΔΔCt values, with Tukey’s multiple comparison test (α = 0.05) to infer significant differences. For animal weight data, a mixed-effects model (restricted maximum likelihood) was used, with Tukey’s multiple comparison test (α = 0.05) to infer significant differences. For animal experiments, a group size of *n* = 5 animals per condition per time point was chosen based on previous studies (*27*). For all data at or below the limit of detection, values were set to the limit of detection.

## Figure Legends

**Figure S1.**
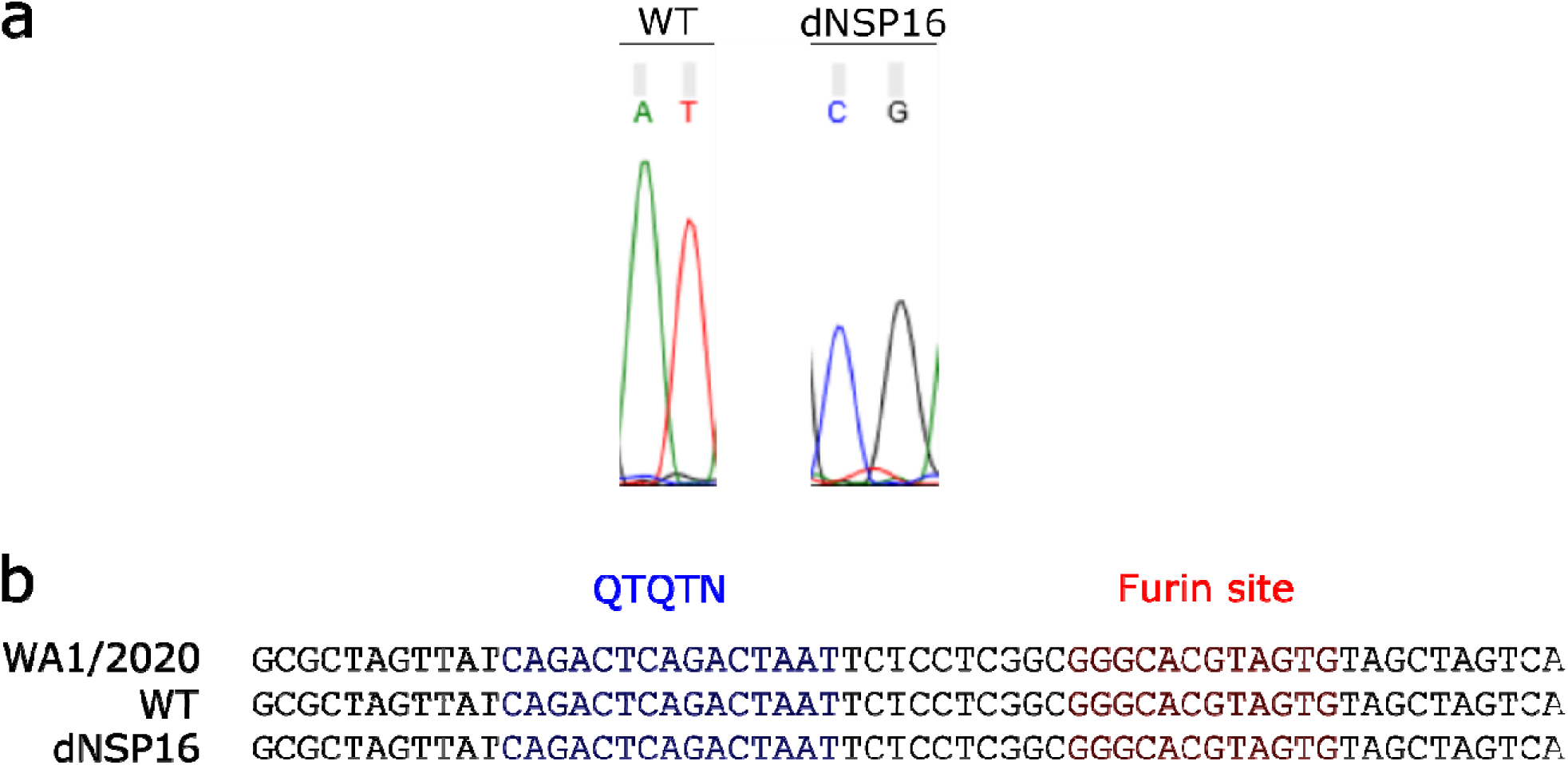
D130 mutation is stable in rescued dNSP16, and rescued infectious clone stocks maintain sequence around furin cleavage site. Viral RNA was extracted from the viral stocks used in the study (“WT” and “dNSP16”). Viral RNA was reverse-transcribed, PCR-amplified around the site of interest, and Sanger sequenced. (a) Shown are the sequencing traces of the 2-base pair site within codon 130 of NSP16 that was mutated from AT to CG to engineer dNSP16. (b) Validated sequence around the furin cleavage site, including the QTQTN motif, for WT and dNSP16, compared to the published sequence for WA1/2020.

**Figure S2.**
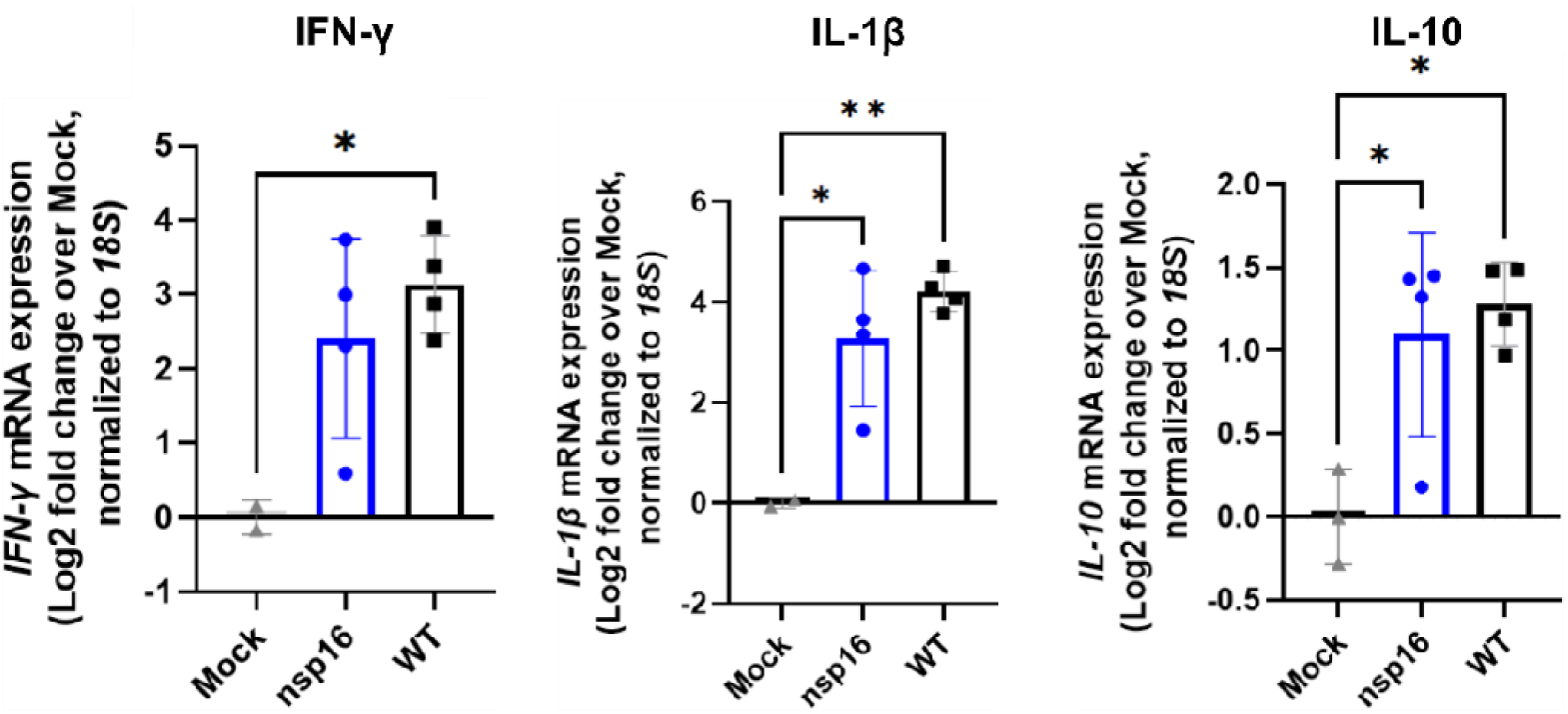
dNSP16 does not drive increased immune gene expression relative to WT. Fold change (log_2_) of expression of the indicated immune genes from lung samples isolated from hamsters infected with the indicated virus (or mock), 2 days post-infection. For each panel, fold changes from dNSP16 or WT samples are measured relative to mock samples. Values from individual hamsters are plotted (symbols) as well as means (bars). Error bars denote standard deviation. All samples were normalized to 18S expression, used as a reference. \**p*<0.05, **p<0.01, \**p*<0.005, \*\*\*\**p*<0.001: results of one-way ANOVA with Tukey’s multiple comparison test (α = 0.05).

**Figure S3.**
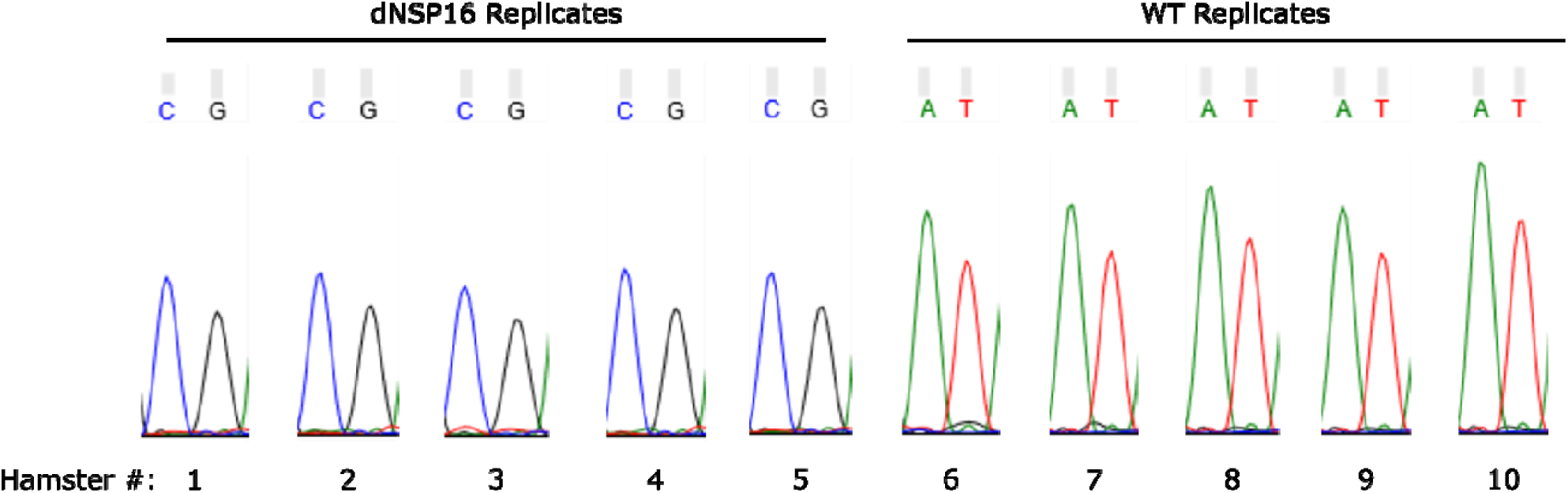
No evidence of reversion of dNSP16 mutation was detected *in vivo*. Viral RNA was extracted from the lungs of hamsters infected with either dNSP16 or WT (numbered 1 through 5 for each group) and which were sacrificed at 4 days post-infection. Viral RNA was reverse-transcribed, PCR-amplified around the site of mutation, and Sanger sequenced. Shown are the sequencing traces of the 2-base pair site within codon 130 of NSP16 that was mutated from AT to CG to engineer dNSP16.

**Figure S4.**
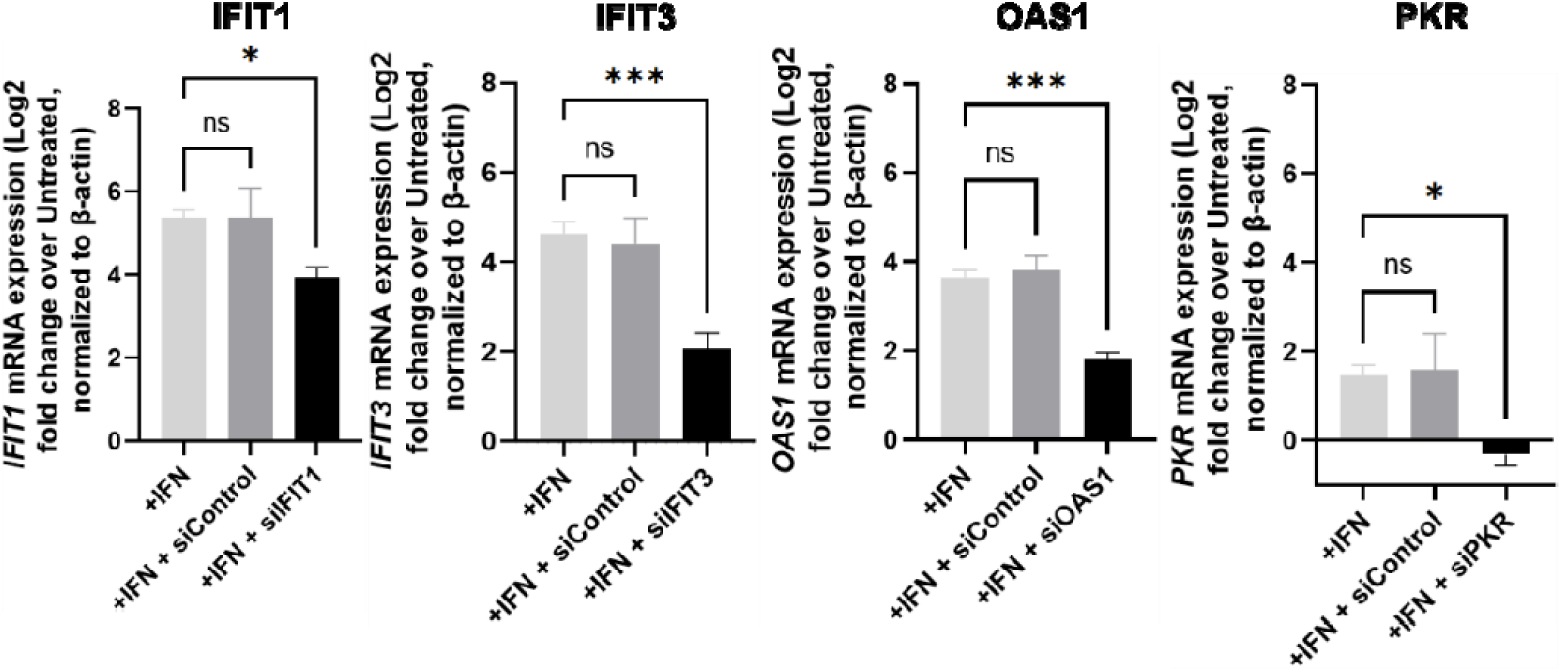
Validation of knockdown of immune gene targets in Vero E6 cells. 1.25 × 10^5^ Vero E6 cells/well were reverse transfected with 1 pmol of the control or gene-specific siRNA 2 days prior to harvest and also treated with 100 U IFN-I one day prior to harvest and assessment of gene expression. Fold change (log_2_) of gene expression is measured relative to untreated samples (i.e. no IFN-I). All samples were normalized to β-actin, used as a reference. \**p*<0.05, \*\*\**p*<0.005, ns = not significant: results of one-way ANOVA with Tukey’s multiple comparison test (α = 0.05). Means are plotted with error bars denoting standard deviation. *n* = 3 biological replicates.

**Figure S5.**
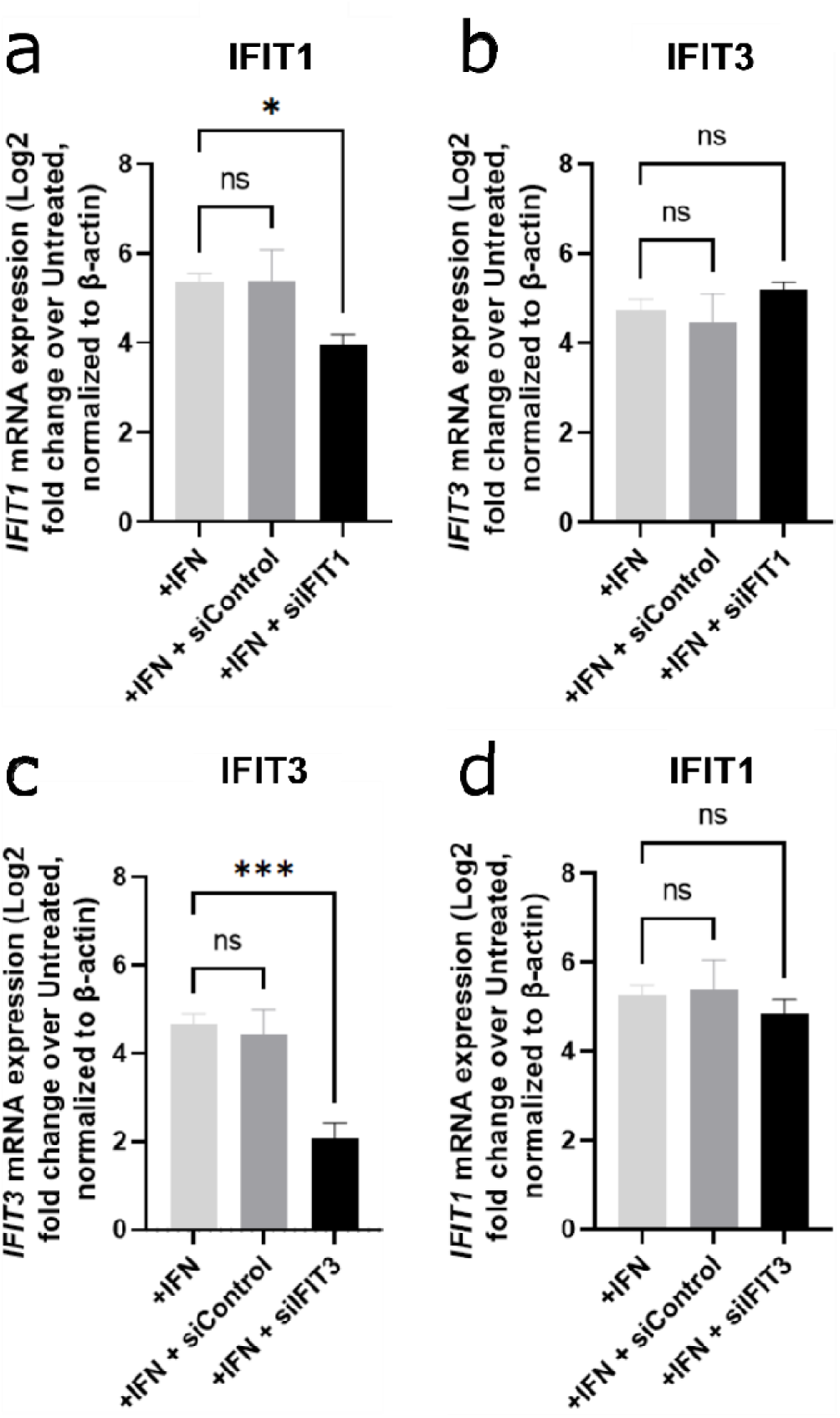
Knockdown of either *IFIT1* or *IFIT3* is specific. 1.25 × 10^5^ Vero E6 cells/well were reverse transfected with 1 pmol/well of either a non-targeting siRNA (“siControl”) or with an *IFIT1*- (a, b) or *IFIT3*- (c, d) targeting siRNA (“siIFIT1” or “siIFIT3”, respectively), or were seeded without treatment. One day later, cells were treated with 100 U of IFN-I to induce interferon-stimulated genes. The following day, cells were lysed for RNA purification and mRNA quantification via reverse transcription and quantitative polymerase chain reaction (PCR). For all panels, gene expression is normalized to β-actin (used as a reference), and fold changes are given relative to untreated controls (i.e. no IFN). \**p*<0.05, \*\*\**p*<0.005, ns = not significant: results of one-way ANOVA with Tukey’s multiple comparison test (α = 0.05). Means are plotted with error bars denoting standard deviation. *n* = 3 biological replicates.

